# Communication as a Two-Person Neural Process: fNIRS Hyperscanning in Adults with and without Post-Stroke Aphasia

**DOI:** 10.64898/2026.09.28.755138

**Authors:** Grace Magee, James Colwell, Shannon Kelley, Yasaa Mohammad, Yiwen Zhang, David Boas, Swathi Kiran, Maria Varkanitsa, Erin Meier, Meryem Yücel

**Affiliations:** Department of Biomedical Engineering, Boston University, Boston, MA; Center for Brain Recovery, Boston University, Boston, MA; Department of Speech, Language, and Hearing Sciences, Boston University, Boston, MA; Department of Communication Sciences and Disorders, Northeastern University, Boston, MA

## Abstract

Communication is inherently interactive, yet most neuroimaging studies examine individuals in isolation. Here, we used functional near-infrared spectroscopy (fNIRS) hyperscanning to investigate interbrain synchrony (IBS) during naturalistic communication. In Study 1, 13 healthy familiar dyads completed Joint Singing, Joint Reading, and Picture Guessing tasks. IBS was quantified using wavelet transform coherence across bilateral frontal and temporoparietal regions. Joint Singing produced higher IBS than the other tasks in several regions, including the dorsolateral prefrontal cortex and temporoparietal junction, and selected effects were greater in true than pseudo-dyads. In Study 2, we applied the same paradigm to five individuals with chronic post-stroke aphasia interacting with trained speech-language pathologists. IBS could be estimated in all dyads after lesion-informed channel exclusion, although regional coverage varied with lesion location. These findings demonstrate that fNIRS hyperscanning can capture task-dependent interpersonal neural coupling during spoken interaction and can be extended to communication involving individuals with aphasia. This approach provides a foundation for studying how neural coordination between communication partners relates to functional communication and partner-supported interaction.

## Introduction

Communication does not occur within isolated individuals, but emerges through dynamic coordination between interacting partners. Successful communication depends on continuous alignment across multiple levels, including timing, rhythm, shared representations, and mutual prediction (Pickering & Garrod, 2004). Despite the inherently interactive nature of communication, much of cognitive neuroscience has relied on single-brain paradigms that reduce communication to isolated stimulus-driven processes (Schilbach et al., 2013). This limitation constrains our ability to study how communication unfolds in real time between people.

Conventional neuroimaging techniques, such as functional magnetic resonance imaging (fMRI), have been critical in identifying the brain regions involved in language production and comprehension (Stephens et al., 2010). However, fMRI requires participants to remain supine and motionless in a confined, noisy scanner environment, making it poorly suited to studying naturalistic, face-to-face communication. To overcome these challenges, functional near-infrared spectroscopy (fNIRS) provides a portable, non-invasive alternative that measures changes in oxygenated and deoxygenated hemoglobin and can be used during face-to-face interaction. fNIRS is well-suited to capturing cortical activity in surface-level language and social brain regions during naturalistic, task-based communication (Yücel et al., 2021; von Lühmann et al., 2021).

Recent advances in two-person neuroscience have built on these methodological developments by introducing hyperscanning, which enables the simultaneous recording of neural activity from multiple interacting individuals (Schilbach et al., 2013). When combined with fNIRS, hyperscanning enables the study of communication as a two-person neural process under ecologically valid conditions (von Lühmann et al., 2021; Yücel et al., 2021). Within this framework, interbrain synchrony (IBS), the alignment or coupling of neural signals across individuals, has been proposed as a neural marker of interpersonal coordination during social interaction (Shamay-Tsoory et al., 2019).

Given the relative newness of hyperscanning methodology, neurobiological models of two-person communication are still in the early stages of development. Nonetheless, systematic reviews of hyperscanning studies to date suggest that intersubject communication, such as turn-taking speech coordination, knowledge-sharing, cooperative creativity, problem-solving, and naturalistic discussion, reliably coincides with significant IBS between interacting partners in the bilateral prefrontal cortex (PFC) and temporoparietal junction (TPJ) (Czeszumski et al., 2022; Kelsen et al., 2022). Gvirts and Perlmutter (2020) proposed that these regions comprise a “mutual social attention system,” while Shamay-Tsoory et al. (2019) advanced a broader “observation-executive” system, including the inferior frontal gyrus (IFG), inferior parietal lobule (IPL), superior temporal sulcus (STS), and premotor cortex, which they described as the core alignment network.

Theories of joint action and dialogue conceptualize communication as an emergent process involving ongoing alignment between partners (Pickering & Garrod, 2004). This alignment can take different forms, including entrainment, or the temporal coordination of behavior, and conceptual alignment, through which partners establish shared meaning and common ground (Pickering & Garrod, 2004). These mechanisms have historically been difficult to investigate at the neural level during real-time interaction (Schilbach et al., 2013). Importantly, hyperscanning evidence suggests that IBS is sensitive to the structure of interaction during communication (Czeszumski et al., 2022). Tasks emphasizing rhythmic or temporal coordination, such as synchronized speech or joint action, tend to elicit strong neural coupling (Osaka et al., 2015), whereas more naturalistic communication tasks engage higher-order cognitive and social processes related to shared understanding (Wang et al., 2022). IBS also appears to reflect interaction-dependent coordination rather than simply similarity in responses to a shared task, as greater synchrony has been reported during real interaction compared with non-interactive or control pairings (Czeszumski et al., 2022). Together, these findings suggest that IBS may provide a means of examining how interpersonal coupling varies across forms of communication that differ in the degree and type of alignment they require.

In addition to task structure, IBS is shaped by characteristics of the interacting partners and their relationship (Kinreich et al., 2017; Zhao et al., 2024). Stronger IBS has been observed between familiar partners, including friends, romantic partners, and long-term collaborators, than between unfamiliar dyads (Kinreich et al., 2017; Zhao et al., 2024). Partner roles and shared goals can also influence IBS, with greater synchrony reported in contexts requiring cooperation, adaptation, or asymmetric expertise (Ellingsen et al., 2023). These findings suggest that interpersonal neural coupling is shaped not only by task demands but also by the social context in which communication occurs. This may be particularly relevant when one partner adapts communication to the needs of the other, as occurs when trained clinicians interact with individuals with aphasia.

Aphasia provides a particularly relevant clinical context for examining these interaction-dependent mechanisms. Aphasia is an acquired language disorder, most commonly following stroke or traumatic brain injury, that affects spoken and written language production and comprehension (Holland, 1982). Importantly, language ability is not equivalent to communication capacity. Audrey Holland’s seminal work demonstrated that some individuals with substantial language impairment can communicate effectively in daily life despite poor performance on standardized language tests, whereas others with milder aphasia may experience substantial difficulty in everyday communication (Holland, 1982). Communication success therefore depends not only on the language abilities of the individual with aphasia, but also on the interaction between communication partners, including turn-taking, shared understanding, repair, and partner adaptation. Although the distinction between language impairment and functional communication has long been recognized in aphasia research, the neural dynamics of communication between individuals with aphasia and their communication partners remain poorly understood. A two-person neuroscience approach may therefore provide an interaction-sensitive means of examining how partner-supported communication unfolds at the neural level.

Despite growing evidence that IBS varies with interaction structure and partner characteristics, important gaps remain. Few hyperscanning studies have directly compared multiple forms of spoken interaction within the same dyads, limiting understanding of how interpersonal coupling changes across different communicative contexts. Moreover, two-person neural approaches have rarely been extended to acquired language impairment, where successful communication may depend strongly on adaptation between partners. Whether fNIRS hyperscanning can provide usable measures of interpersonal coupling in individuals with focal cortical lesions therefore remains unclear.

The present study used fNIRS hyperscanning to examine IBS during interactive communication in two complementary studies. Study 1 examined neurologically healthy adults interacting with familiar partners, whereas Study 2 extended the same paradigm to individuals with chronic post-stroke aphasia interacting with trained speech-language pathologists. Across both studies, dyads completed three interactive tasks selected to sample communication contexts with different interaction structures. Joint Singing provides strong shared rhythmic, melodic, and temporal structure, creating continuous opportunities for interpersonal entrainment. Joint Reading also requires simultaneous vocal production and coordination of pacing and prosody, but without the same musical scaffold. Picture Guessing is less temporally constrained and instead requires alternating speaker-listener roles, description, inference, clarification, repair, and convergence on shared meaning. Thus, rather than isolating a single communicative mechanism, the three tasks provide complementary contexts that differ in the degree and form of interpersonal coordination they require.

The present study had two aims. First, in healthy familiar dyads, we tested whether IBS varied across Joint Singing, Joint Reading, and Picture Guessing within predefined frontal and temporoparietal ROIs. We hypothesized that IBS magnitude would differ across tasks, with the strongest synchrony expected during Joint Singing because of its highly constrained rhythmic and temporal structure. Second, we evaluated the feasibility of applying the same fNIRS hyperscanning paradigm to individuals with chronic post-stroke aphasia interacting with trained speech-language pathologists. Specifically, we asked whether usable IBS estimates could be obtained during these communication tasks after accounting for channels overlapping lesioned cortex. Given the small sample size and heterogeneity in lesion distribution and aphasia profile, analyses of the aphasia-SLP dyads were considered exploratory and descriptive.

By first examining task-sensitive IBS in healthy familiar dyads and then extending the same paradigm to aphasia-SLP dyads, this work links foundational questions in two-person neuroscience with a clinically important model of disrupted communication after stroke.

## Methods

### Probe Design and Regions of Interest

Five bilateral cortical regions of interest were selected a priori based on their involvement in language and social-cognitive/interpersonal processing [Gan et al., 2024; Merchant et al., 2025; Schurz et al., 2021]: the inferior frontal gyrus (IFG), dorsolateral prefrontal cortex (dlPFC), temporoparietal junction (TPJ), supramarginal gyrus (SMG), and middle frontal gyrus (MFG). A fNIRS probe was designed in AtlasViewer (Aasted et al., 2015) using standard international 10-20 EEG coordinates and a three-dimensional adult head model (Holmes et al., 1998). Source and detector locations were selected to maximize coverage of these cortical regions while incorporating short-separation channels to regress superficial physiological signals. Because the probe was constructed on a low-density cap with a limited number of source-detector pairs, channels corresponding to each ROI were selected based on the greatest spatial overlap between their measurement sensitivity profiles and the anatomically defined cortical regions.

The final probe was implemented on a whole-head fNIRS cap with 16 dual-wavelength sources at 760 and 850 nm, and 16 detectors. The probe included both long-separation channels, approximately 30 mm source-detector distance, and short-separation channels, approximately 8 mm source-detector distance. The final configuration included 41 channels, consisting of 33 long-separation channels distributed across frontal, temporal, and parietal regions and 8 short-separation channels. The same probe configuration was used for all participants. A schematic of the probe layout is shown in Figure 1.

**Figure 1.**
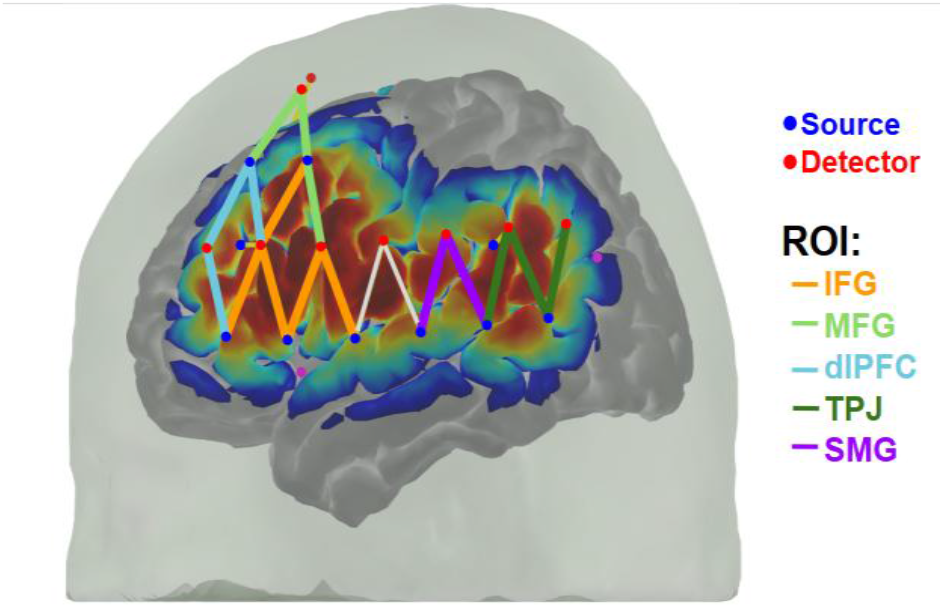
fNIRS hyperscanning probe layout. Probe configuration used for hyperscanning data acquisition, illustrating the placement of sources, detectors, and measurement channels over the bilateral IFG, DLPFC, MFG, TPJ, SMG.

### Data Acquisition and Synchronization

Each participant wore an identical whole-head fNIRS cap populated with the same source-detector array. Data were acquired from each member of the dyad using independent NIRSport2 systems operated in hyperscanning mode. Cap size was selected based on head circumference measurements obtained before the study visit. At the start of the session, study team members situated the cap on the participant’s head to ensure proper positioning of the cap markers relative to participant anatomy (e.g., Cz, pre-auricular points).

Before recording, optode-scalp coupling was optimized by parting the hair beneath each optode and adjusting optode placement to improve contact and reduce light attenuation. Signal quality was monitored in real time using the Aurora acquisition software signal quality map. Optodes were adjusted until the best achievable signal quality was obtained across the probe array.

The two fNIRS systems were synchronized using the extension port and wireless communication between systems. Experimental triggers were delivered simultaneously to both systems using Lab Streaming Layer. Behavioral tasks were implemented in PsychoPy and synchronized with Aurora/Hyperscan software to ensure temporal alignment between the task events and fNIRS recordings. Data were sampled at 23.3 Hz, the highest sampling rate supported by the acquisition configuration used in this study.

### Experimental Tasks

Dyads completed three interactive communication tasks: Joint Singing, Joint Reading, and Picture Guessing. Each active task block was preceded and followed by a 30-second rest period. During rest periods, participants were instructed to remain still and keep their eyes closed to minimize visual interaction and reduce visually driven synchrony.

In Joint Singing, dyads sang a familiar melody together during two 120-second blocks. In Joint Reading, dyads read a shared narrative aloud during two 120-second blocks. In Picture Guessing, one partner described a common object or picture while the other partner guessed its identity; roles alternated across trials during two 300-second blocks. These tasks were selected to vary in communicative structure, rhythmic coordination, shared speech production, and interactive meaning exchange while allowing multiple hemodynamic response cycles to be observed. Participants were instructed to minimize head and body movement during all tasks. A schematic of the task design is shown in Figure 2.

**Figure 2.**
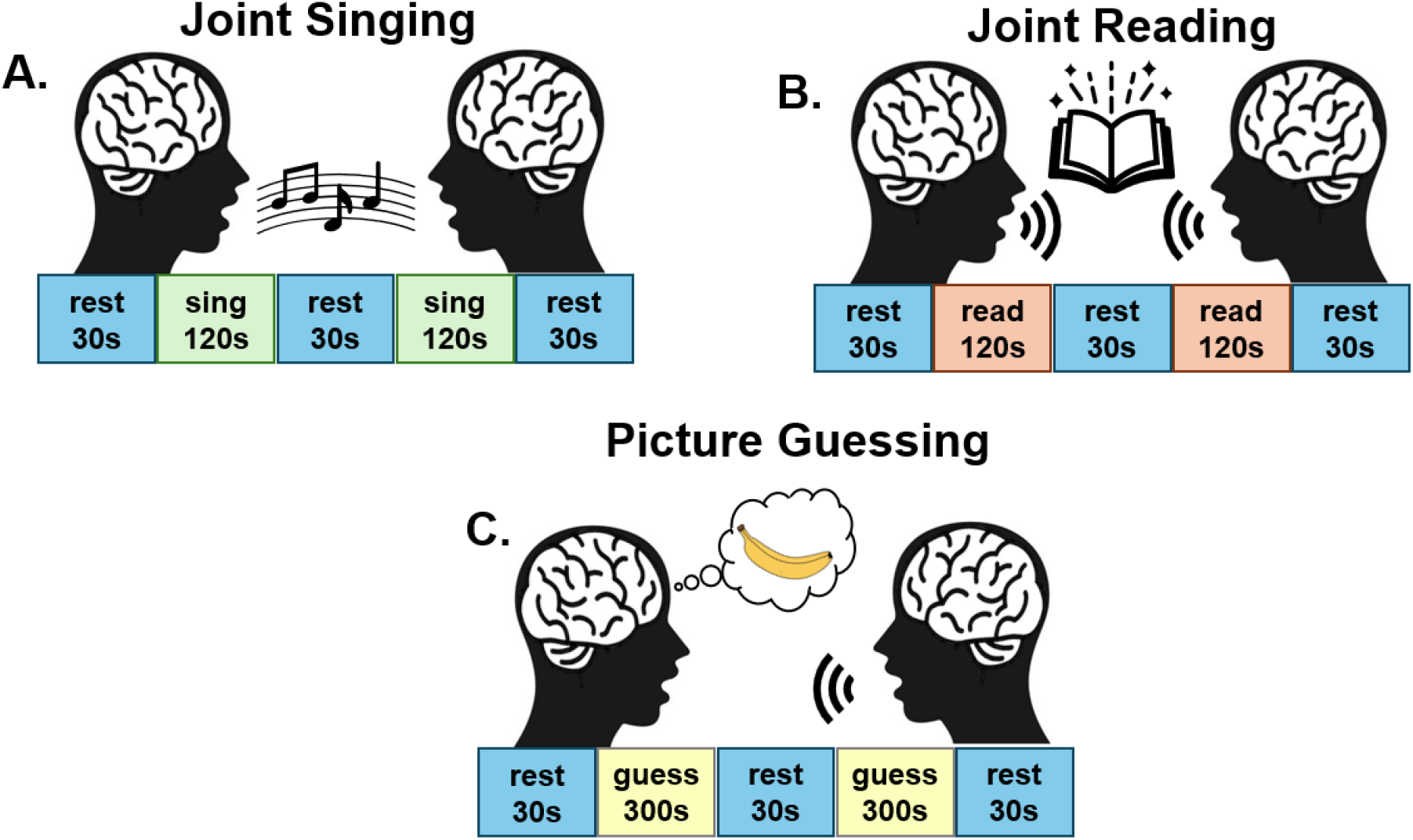
Overview of the three interactive communication tasks completed by each dyad: **(A)** Joint Singing, **(B)** Joint Reading, and **(C)** Picture Guessing. Joint Singing and Joint Reading each comprised two 120-s task blocks, whereas Picture Guessing comprised two 300-s task blocks with alternating speaker and listener roles. All tasks included 30-s baseline rest periods before, between, and after task blocks.

### fNIRS Preprocessing and Hemodynamic Response Estimation

Data preprocessing was performed in MATLAB using a modified dual-subject hyperscanning pipeline based on Homer3 (Huppert et al., 2009). Raw light intensity data were converted to optical density. Channels with a signal-to-noise ratio below 3 were pruned and excluded from subsequent analyses. Motion artifacts were corrected using wavelet filtering (Molavi & Dumont, 2012) with an interquartile range threshold of 1.5. To preserve slow task-related hemodynamic fluctuations during the long task blocks, no high-pass filter or band-pass filter was applied. Instead, slow baseline drift was modeled with a first-order polynomial, and high-frequency noise was attenuated with a low-pass filter at 0.5 Hz. Optical density signals were then converted to changes in oxygenated hemoglobin, HbO, and deoxygenated hemoglobin, HbR, using the modified Beer-Lambert law, with no differential pathlength correction.

Task-evoked hemodynamic responses were estimated using a general linear model (GLM). The hemodynamic response was modeled using a consecutive sequence of Gaussian basis functions, with σ = 1.0 s and 1.0 s spacing, to allow flexible estimation of the response over the extended task blocks. Short-separation regression was included in the GLM to reduce superficial physiological contributions. For each long-separation channel, the short-separation channel with the highest temporal correlation was selected as the nuisance regressor (Gagnon et al., 2014).

The modeled response window included the full task block and an additional 15 s after task offset to capture the return toward baseline.

### Region of Interest-level Signal Extraction

Channel locations were mapped to standard anatomical space using AtlasViewer (Aasted et al., 2015). Source-detector channel coordinates were registered to the Colin27 head model (Holmes et al., 1998) and projected to Montreal Neurological Institute space and assigned to anatomical regions using BioImage Suite (Papademetris et al., 2006) and Brodmann atlas labels (Lacadie et al., 2008). Channels assigned to the same anatomical region were grouped into region-of-interest averages. ROI-level HbO time series were computed by averaging across all retained long-separation channels assigned to each ROI.

### Interbrain synchrony analysis

Interbrain synchrony was quantified using wavelet transform coherence (WTC) between homologous ROI-level HbO time series from the two members of each dyad. WTC estimates the local correspondence between two signals as a function of both time and frequency, allowing coupling that varies over the course of an interaction to be characterized rather than assuming a stationary relationship across the entire recording. For each dyad, homologous regions were paired across participants; for example, HbO activity from participant A’s left IFG was paired with HbO activity from participant B’s left IFG. Analyses focused on HbO because of its typically larger task-related response and higher signal-to-noise ratio in fNIRS measurements. Analyses were restricted to the 0.08 to 0.30 Hz frequency range, consistent with prior fNIRS hyperscanning studies (Hirsch et al., 2017). Coherence values were averaged within this frequency range and across task blocks to obtain one IBS value per dyad, task, and homologous ROI pair. IBS values range from 0 to 1, with higher values indicating greater temporal correspondence between the partners’ HbO signals within the specified frequency range.

### Study 1 Methods

Study 1 included twenty-six healthy adults forming 13 dyads (14 male, 12 female; mean age 24.5 ± 8.7 years). Study 1 was approved by the Boston University Institutional Review Board under protocol 4502, and Study 2 under protocol 4966E. All participants provided informed consent before participation. Participants reported no history of neurological or psychiatric disorders and were not taking medications known to affect cerebral blood flow. All participants were able to engage in cooperative communication tasks and completed the full experimental protocol. All participants were familiar with their dyad partner. Data acquisition, task implementation, preprocessing, and interbrain synchrony analyses were conducted according to the procedures described in the General Methods section.

The primary outcome for Study 1 was interbrain synchrony, quantified as the mean wavelet transform coherence value for each dyad, task, and homologous ROI pair. This yielded task-specific IBS estimates for Joint Singing, Joint Reading, and Picture Guessing across the predefined bilateral ROIs. The primary analysis examined whether IBS varied across communicative tasks, reflecting differences in the alignment demands imposed by each task.

Statistical analyses were conducted at the dyad level using mean IBS values extracted for each dyad, task, and homologous ROI pair. Because of the sample size and the distribution of coherence values, nonparametric tests were used. Task-related differences in IBS were examined separately for each ROI using pairwise Wilcoxon signed-rank tests comparing Joint Singing, Joint Reading and Picture Guessing within the same dyads. P-values were not adjusted for multiple comparisons using false discovery rate (FDR) correction.

To assess whether observed interbrain synchrony reflected interpersonal interaction rather than shared task timing or chance correlations, pseudo-dyads were generated by re-pairing participants across original dyads. Each pseudo-dyad consisted of two participants from different original dyads who had completed the same experimental tasks. This approach preserved task structure and timing while removing the original communicative interaction between partners.

Interbrain synchrony was calculated for pseudo-dyads using the same analysis pipeline applied to true dyads. Synchrony values from true and pseudo-dyads were then compared by task and region of interest.

### 3. Study 2 Methods

Study 2 included five individuals with chronic post-stroke aphasia (PWA) and five trained speech-language pathologists (SLP), forming five aphasia-SLP dyads. All PWA were recruited through the Boston University Center for Brain Recovery. All PWA had a documented unilateral cerebrovascular accident affecting language-relevant cortical regions (see lesion overlay for four PWA with available stroke lesion maps in Figure 3), were 61–73 years old (M = 66.8, SD = 5.3), and had 16–20 years of education (M = 17.8, SD = 1.5). Time post-onset ranged from 60 to 565 months (M = 196.8, SD = 210.7), indicating a chronic sample with substantial variability in time since stroke. Aphasia severity was mild overall, with WAB-R Aphasia Quotient scores ranging from 85.2 to 96.4 (M = 92.3, SD = 4.6). PWA who scored above the WAB-R Aphasia Quotient cutoff endorsed persistent language challenges that differed from their pre-stroke baseline. WAB-R Reading scores ranged from 84 to 100 (M = 94.4, SD = 6.2). They were required to be medically stable and able to speak and read. Exclusion criteria included a history of additional neurological disorders, severe cognitive impairment precluding task participation, or contraindications to fNIRS data acquisition. The SLP partners were 27–37 years old (M = 31.4, SD = 3.6) and had 18–25 years of education (M = 20.8, SD = 2.9). For subject-level demographic and clinical characteristics, see Table 1.

**Table 1.** Demographic and clinical characteristics of participants with aphasia (PWA).

| Participant | Dyad | Age | Education | MPO | WAB-R AQ | WAB-R Reading |
| --- | --- | --- | --- | --- | --- | --- |
| PWA01 | 1 | 71 | 16 | 85 | 96.3 | 96 |
| PWA02 | 2 | 73 | 18 | 60 | 92.1 | 94 |
| PWA03 | 3 | 67 | 20 | 94 | 85.2 | 84 |
| PWA04 | 4 | 61 | 17 | 565 | 91.3 | 98 |
| PWA05 | 5 | 62 | 18 | 180 | 96.4 | 100 |
| SLP01 | 1 | 31 | 25 | NA | NA | NA |
| SLP02 | 2 | 31 | 18 | NA | NA | NA |
| SLP03 | 3 | 31 | 22 | NA | NA | NA |
| SLP04 | 4 | 37 | 21 | NA | NA | NA |
| SLP05 | 5 | 27 | 18 | NA | NA | NA |

**Figure 3.**
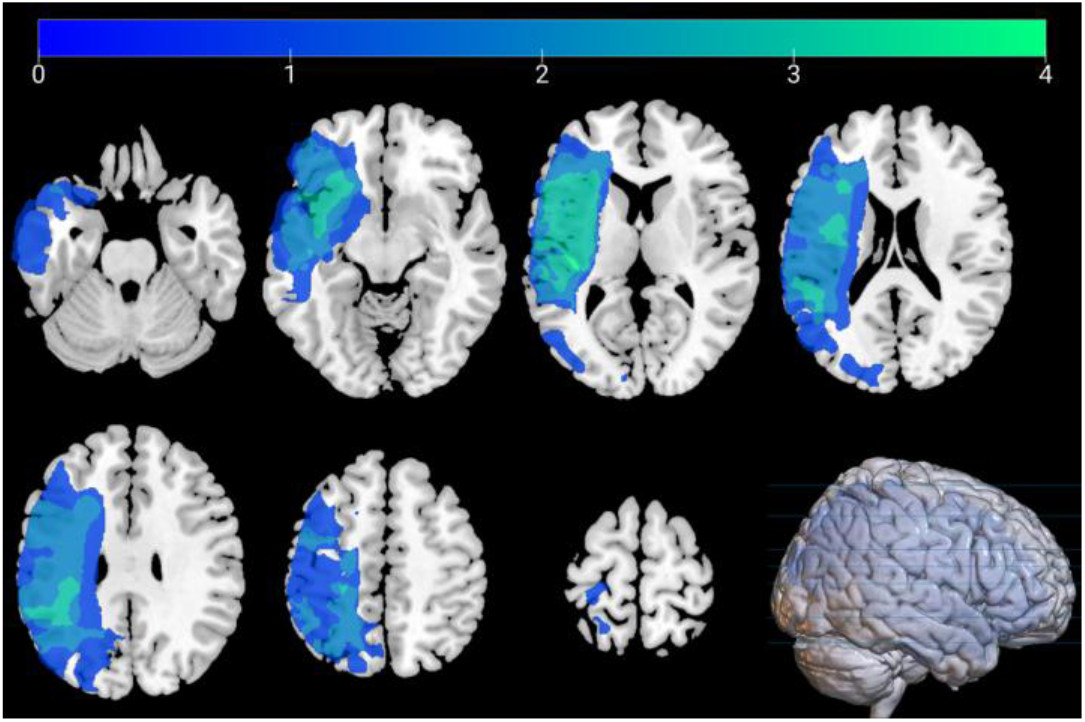
Lesion overlay for four PWA with available stroke lesion maps.

All PWA-SLP dyads completed the experimental tasks and data acquisition procedures described in the General Methods section, using the same standardized hyperscanning protocol. For Study 2, task instructions were repeated or simplified as needed, and practice trials were completed before hyperscanning to ensure that participants understood the task requirements.

To reduce contributions from long-separation channels whose nominal cortical sampling regions substantially overlapped lesioned tissue, lesion-informed channel exclusion was performed before applying the ROI-level analysis. Channel-level lesion status was determined using an MNI-registered channel map and subject-specific 3D lesion reconstructions. For each channel, a spherical region of interest (radius = 6 mm) was generated using the channel’s MNI coordinates as the sphere’s center. Each participant’s 3D lesion mask was then overlaid with the channel-centered spheres to evaluate spatial overlap between lesion tissue and each channel’s cortical sampling region.

Channels were labeled as “intact,” “intact (perilesional),” or “lesioned.” Channels were classified as *intact (perilesional)* when the sphere region was predominantly intact (operationally defined as >80% intact tissue). Channels were classified as *lesioned* when the sphere region was fully lesioned or showed >20% overlap with lesioned tissue. Only channels labeled “lesioned” were excluded from subsequent ROI-level analyses. Channels labeled “intact” or “intact (perilesional)” were retained for analysis. Short-separation channels were not subjected to lesion-based exclusion and were retained for superficial signal regression.

Given the small sample size, Study 2 was analyzed descriptively rather than with formal inferential statistics. For each PWA–SLP dyad, IBS values were calculated for each task and retained homologous ROI pair using the procedures described in the Shared fNIRS Hyperscanning Protocol. To derive a summary measure of dyadic synchrony, ROI-level IBS values were averaged across the retained bilateral ROIs, yielding a single network-level IBS value for each dyad and task. These network-level values were used to visualize task-related patterns of IBS across PWA–SLP dyads.

## Results

Mean IBS values were compared across Joint Singing, Joint Reading, and Picture Guessing for each bilateral ROI (Figure 4). Joint Singing showed numerically higher IBS than Joint Reading and Picture Guessing across several frontal and temporoparietal regions.

**Figure 4.**
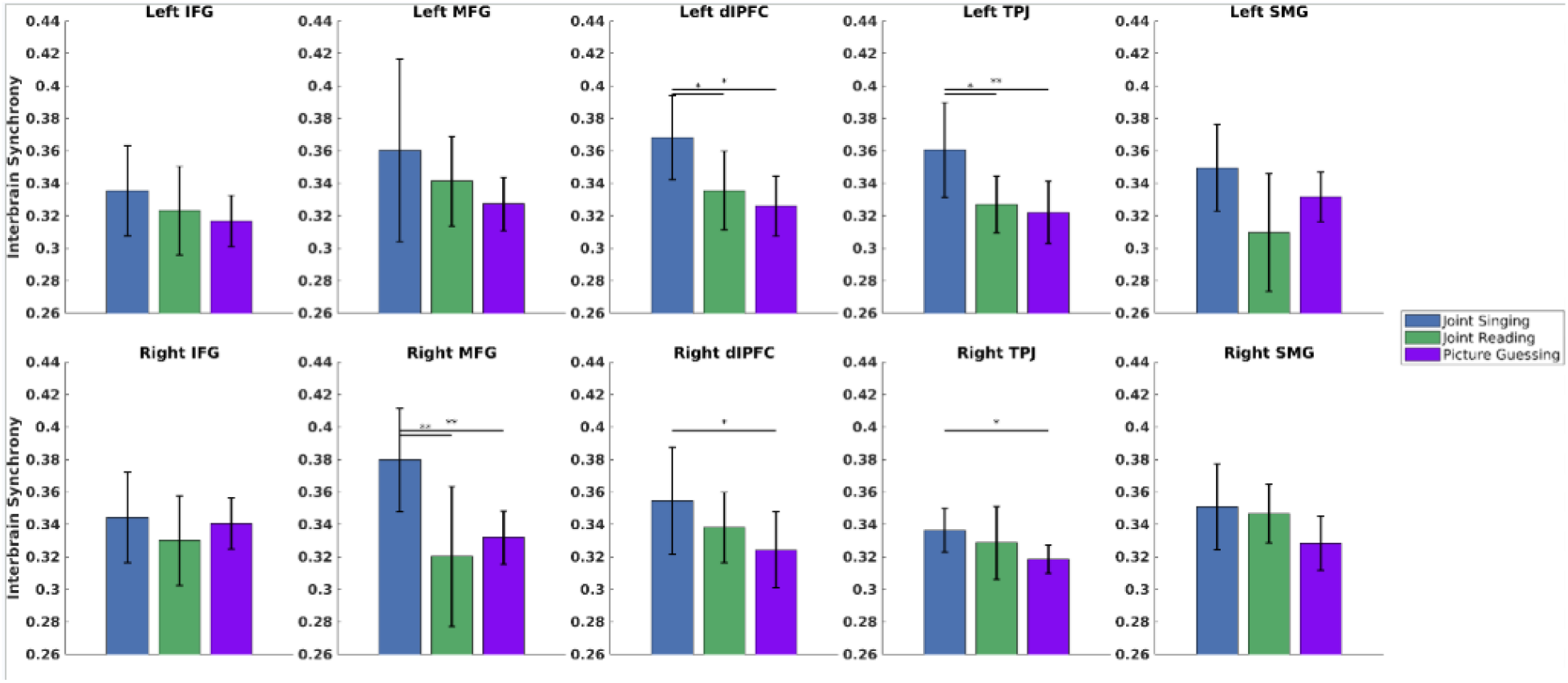
Mean IBS across bilateral ROIs during Joint Singing, Joint Reading, and Picture Guessing. Bars represent mean IBS ± standard deviation. Asterisks indicate significant pairwise comparisons (*p < 0.05, **p < 0.01, *** p < 0.001)

Pairwise Wilcoxon signed-rank tests revealed significant task-related differences in several frontal and temporoparietal ROIs. In the left dlPFC, Joint Singing elicited significantly greater IBS than Joint Reading (*p* = 0.02) and Picture Guessing (*p* = 0.02). Similarly, in the left TPJ, IBS during Joint Singing was significantly higher than during Joint Reading (*p* = 0.04) and Picture Guessing (*p* = 0.009). In the right dlPFC, Joint Singing also resulted in significantly greater IBS than Picture Guessing (*p* = 0.04). Within the right MFG, Joint Singing demonstrated significantly greater IBS than both Joint Reading (*p* = 0.002) and Picture Guessing (*p* = 0.002). Finally, the right TPJ exhibited significantly greater IBS during Joint Singing compared with Picture Guessing (*p* = 0.02).

No statistically significant task-related differences were observed in the bilateral IFG, left MFG and bilateral SMG beyond the comparisons made in Figure 4. Overall, these findings indicate that Joint Singing was associated with selectively higher IBS in bilateral frontotemporoparietal regions, particularly the DLPFC, TPJ, and right MFG, rather than producing a uniform increase in synchrony across all ROIs. Comparison of pseudo and true dyads revealed significantly greater IBS in the true dyads within the left dlPFC during Joint Singing (*p* = 0.03) and the left TPJ during Joint Singing (*p* = 0.05) (Figure 5). No other left-hemisphere ROIs showed significant differences between pseudo and true dyads across Joint Singing, Joint Reading, or Picture Guessing.

**Figure 5.**
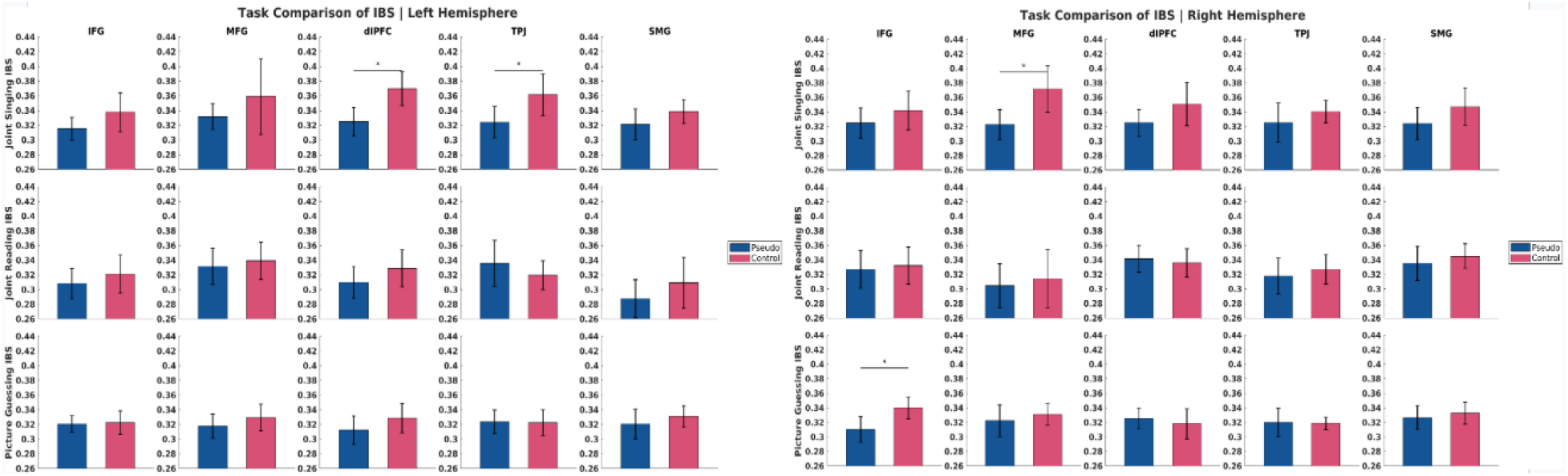
Comparison of interbrain synchrony (IBS) between true dyads and pseudo-dyads across experimental tasks in the left hemisphere (left) and the right hemisphere (right). Mean IBS values are shown for true dyads (red) and pseudo-dyads (blue) within the five ROIs. Asterisks denote statistically significant differences between true and pseudo-dyads (p < 0.05, uncorrected).

Within the right hemisphere, true dyads exhibited significantly greater IBS than pseudo-dyads in the right MFG during Joint Singing (*p* = 0.02) and the right IFG during Picture Guessing (*p* = 0.02) (Figure 5). No other right hemisphere ROIs showed significant differences between pseudo and true dyads across Joint Singing, Joint Reading, or Picture Guessing.

IBS could be estimated in all five aphasia-SLP dyads following lesion-informed channel exclusion, although retained ROI coverage varied across participants according to lesion location and available channels. At the network level, mean IBS was descriptively highest during Joint Singing, followed by Joint Reading and Picture Guessing (Figure 6), with substantial variability across dyads. ROI-level IBS values following lesion-informed channel exclusion are shown descriptively in Figure 7. Given the small sample size, no inferential comparisons between tasks were performed.

**Figure 6.**
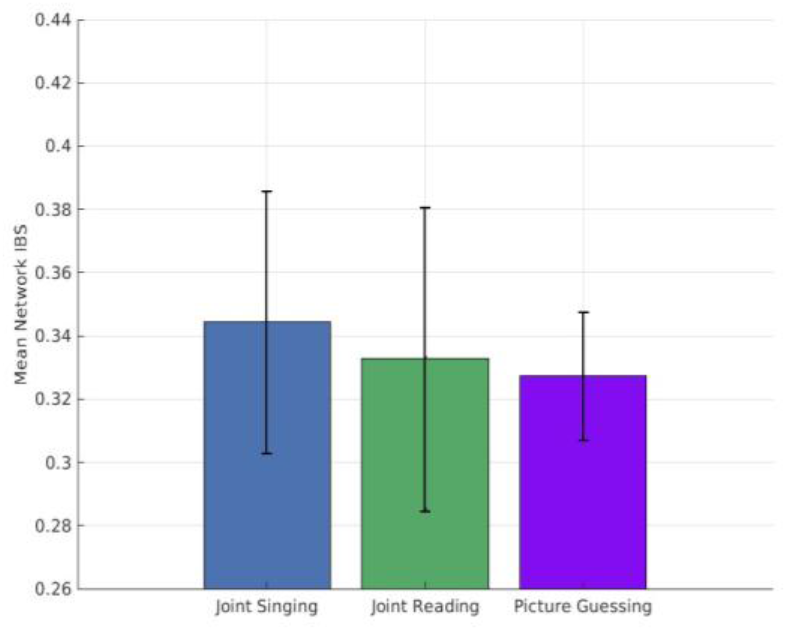
Mean network IBS across study 2 dyads during Joint Singing, Joint Reading, and Picture Guessing. Bars represent mean IBS ± standard deviation.

**Figure 7.**
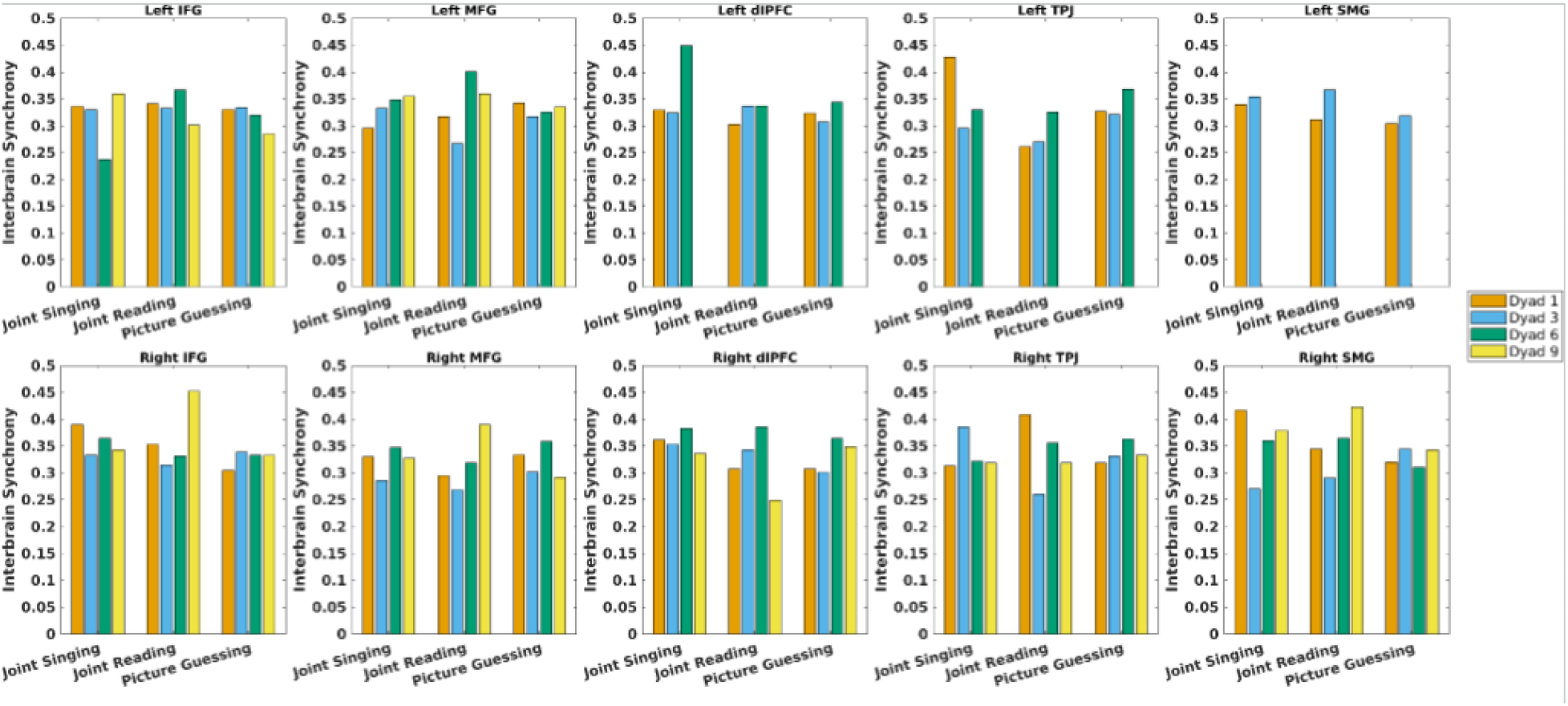
Mean interbrain synchrony (IBS) across bilateral regions of interest (ROIs) during Joint Singing, Joint Reading, and Picture Guessing after exclusion of lesion-affected channels in Study 2 participants with aphasia. Channels overlying lesioned cortical tissue were excluded prior to analysis. Depending on lesion location and extent, exclusion of these channels resulted in insufficient coverage for analysis in specific ROIs for some dyads.

## Discussion

The present study used fNIRS hyperscanning to examine IBS during interactive communication in two complementary studies. Study 1 tested whether IBS varies as a function of communicative task demands in healthy familiar dyads. Study 2 extended the same paradigm to individuals with chronic post-stroke aphasia interacting with trained SLPs. Together, the studies show that fNIRS hyperscanning can capture task-dependent variation in IBS in healthy familiar dyads and can be feasibly extended to communication involving individuals with chronic post-stroke aphasia.

Although preliminary, several findings from Study 1 provide insight into the nature of interpersonal neural synchrony across different types of language-based interaction. A central challenge in hyperscanning research is to establish mechanistic accounts of IBS, beginning with evidence that observed coupling is not merely epiphenomenal but reflects features of genuine interpersonal interaction. A true-versus-pseudo dyad difference would provide evidence that at least part of the observed synchrony cannot be explained solely by shared task timing or coincident temporal structure.

To our knowledge, no prior study has directly compared IBS between tasks requiring speech entrainment or mutual understanding within the same sample of dyads. Nonetheless, high IBS for Joint Singing aligns with other hyperscanning studies that report significant IBS for tasks involving rhythmic, motoric, or vocal coordination and entrainment between interlocutors (e.g., İşbilir, 2016; Kawasaki et al., 2013; Osaka et al., 2015; Pérez et al., 2017). For example, in an experiment that closely mirrors our Joint Singing task, Osaka et al. (2015) asked 15 young, healthy pairs to sing popular Japanese nursery rhymes in different conditions. They found significantly greater neural coherence between pairs for joint compared to independent singing in the left inferior frontal cortex and right middle temporal gyrus, effects that were similar when pairs were facing each other or a barrier. In Kawasaki et al. (2013), 40 young, healthy adults alternated producing the alphabet with either another human or a robot while undergoing EEG. Significant cross-correlations in EEG frequency amplitudes—indicative of interpersonal neural synchrony—were observed in the left inferior frontal, left premotor, and bilateral temporoparietal cortex during human-human (but not human-robot) interactions. Temporoparietal activations, which approximated TPJ, were significantly correlated with synchrony of inter-partner alignment of speech durations and intervals.

At a regional level, significant task effects in Study 1 were observed in the bilateral dlPFC and TPJ, regions proposed to serve as core hubs within the relational neuroscience network (De Felice et al., 2025; Gvirts & Perlmutter, 2020). Indeed, in their meta-analysis of fNIRS hyperscanning studies, Czeszumski et al. (2022) found that PFC was implicated across all 13 included studies, while significant TPJ effects were reported in four investigations. They also reported a large effect size when contrasting cooperation and control conditions, with many entrainment-type paradigms among the included studies. The systematic review by Kelsen et al. (2022) concluded that IBS during spoken communication occurred predominantly in frontal and temporoparietal regions, and their study-by-study synthesis repeatedly highlighted dlPFC and TPJ in cooperative/creative communication studies. In larger and more recent meta-analyses, Lotter et al. (2023) and Berni et al. (2026) reported convergent results in right TPJ across hyperscanning experiments. In Lotter et al. (2023), their follow-up meta-analytic coactivation analysis showed that right TPJ is embedded in a network including bilateral TPJ and bilateral dlPFC in addition to more medial and subcortical regions.

Thus, our regional results from Study 1 are consistent with the extant literature. Within the social cognition literature, dlPFC is thought to contribute to perspective-taking and the executive demands of real-time interpersonal coordination, whereas TPJ is implicated in mentalizing and reciprocal social interaction (Frith & Frith, 2006; Gan et al., 2024; Healey & Grossman, 2018; Merchant et al., 2025; Van Overwalle, 2009). The stronger IBS effects in these regions during Joint Singing may reflect the continuous and structured coordination of this task. Joint singing requires ongoing prediction, monitoring, and adjustment of one’s vocal timing relative to a partner (Delius & Müller, 2023; Keller et al., 2014), with shared rhythmic and melodic cues providing a strong scaffold for maintaining alignment. Although Joint Reading likewise requires continuous interpersonal coordination, it lacks these additional rhythmic and melodic cues to maintain constant alignment. In contrast, the demands for perspective-task and online coordination during Picture Guessing likely vary across each trial as partners progressively converge on a shared conceptual representation. These interpretations remain speculative, however, because Study 1 did not include behavioral measures of interpersonal alignment. Therefore, future work should examine whether task-related differences in regional IBS correspond to differences in speech synchrony and/or conceptual alignment between partners.

Importantly, prior literature has demonstrated that IBS is not simply an entrainment phenomenon. A recent meta-analytic study by Jiang et al. (2026) reported that a substantial overall EEG-based IBS effect persisted after explicitly removing motor and rhythm-based paradigms. Individual studies have also shown significant interpersonal neural coupling while partners establish shared meanings and communication systems (Aili et al., 2025; Chen et al., 2022; Domínguez-Arriola et al., 2026; Hirsch et al., 2018; Jiang et al., 2012; J. Liu et al., 2023; Y. Liu et al., 2017; Ono et al., 2022; Pérez et al., 2017; Speer et al., 2024; Stolk et al., 2014). For example, by explicitly modeling the speech envelopment during oral narratives, Pérez et al. (2017) found that speaker-listener synchronization in brain oscillations was mediated by low-level sensory mechanisms but also broader situational interactive processes. Stolk et al. (2014) used fMRI hyperscanning during a communicative game in which one partner conveyed a target token configuration to another. Interbrain coupling in right superior temporal gyrus was greater for novel than established communicative problems, was specific to real dyads, and occurred at a timescale corresponding to problem-solving episodes rather than sensorimotor events. These and related findings indicate that IBS can reflect higher-order interpersonal processes involved in establishing shared meaning and conceptual alignment.

Study 2 provides a preliminary extension of the paradigm to communication involving individuals with chronic post-stroke aphasia. IBS could be quantified after lesion-informed channel exclusion in all five aphasia-SLP dyads, demonstrating the feasibility of deriving dyadic fNIRS measures during these communication tasks despite heterogeneous lesion distributions.

Descriptively, network-level IBS was highest during Joint Singing, followed by Joint Reading and Picture Guessing, the same ordering observed at the group level in Study 1. Given the very small sample, however, this pattern should not be interpreted as evidence that the task effects observed in Study 1 replicate in aphasia. Rather, it provides an initial observation that can be tested in a larger sample. Importantly, lower IBS should not be equated with poorer communication. Joint Reading and especially Picture Guessing involve less temporally constrained interaction, in which successful communication may depend on flexible turn-taking, repair, inference, and partner accommodation rather than continuous time-locked synchrony. Future work incorporating behavioral measures of these processes will be necessary to determine how different forms of interpersonal coupling relate to communicative success in aphasia.

Study 2 motivates several directions for future research. First, larger samples are needed to test task effects in aphasia–SLP dyads statistically and to determine whether the patterns observed here generalize across aphasia severity, lesion profiles, and communication contexts. Second, future studies should integrate behavioral measures of dyadic alignment, including acoustic entrainment, speech timing, turn-taking, repair sequences, naming success, discourse alignment, communicative effectiveness, and subjective ratings of partner experience. These measures will be essential for determining whether variation in IBS relates to meaningful communicative success. Third, future work should examine how partner characteristics shape IBS, including communication partner training, familiarity, therapeutic alliance, and accommodation capacity. These factors are clinically important because the burden of successful communication in aphasia is shared between the person with aphasia and the communication partner.

### Limitations

Several limitations should be noted. Study 1 included a modest sample of healthy young adults who were familiar with their partners, and Study 2 included only five aphasia–SLP dyads. Therefore, the present findings should be interpreted as pilot data rather than definitive evidence of task-or group-level effects. The two studies were complementary but not designed as matched group comparisons. In addition, the aphasia sample was relatively mild overall, and lesion reconstructions were available for only a subset of participants. The current analyses focused on homologous ROI-to-ROI coherence and averaged IBS across relatively long task blocks; therefore, they may not capture time-lagged, cross-regional, or event-level coupling that may be especially important during flexible communication tasks such as Picture Guessing. Finally, behavioral interaction measures were not yet integrated with IBS, limiting our ability to link neural synchrony directly to communicative success.

Because fNIRS signals contain both cortical and systemic hemodynamic components, and because Joint Singing may synchronize respiration, vocal effort, and movement between partners, greater HbO coherence during this condition cannot be attributed exclusively to neural coupling. Short-separation regression was used to reduce superficial physiological contributions, but future studies incorporating concurrent systemic physiological measurements will be important for distinguishing cortical coupling from shared peripheral physiology. In addition, lesion-informed channel exclusion was based on the spatial overlap between lesion masks and coordinate-centered spherical regions rather than participant-specific photon-sensitivity profiles; therefore, lesion overlap should be interpreted as an operational estimate of whether a channel sampled lesioned tissue.

In summary, these studies demonstrate that fNIRS hyperscanning can characterize interpersonal HbO synchrony across different forms of spoken interaction and can be extended to communication involving individuals with chronic post-stroke aphasia. In healthy familiar dyads, IBS varied across communicative tasks, with greater synchrony during the highly temporally structured Joint Singing condition in several frontal and temporoparietal regions. In aphasia-SLP dyads, lesion-informed processing enabled IBS estimation across heterogeneous lesion profiles, establishing feasibility for larger clinical studies. Linking these neural measures with behavioral indices of coordination, accommodation, and communicative success will be critical for determining the functional significance of interpersonal synchrony in everyday communication.

## Acknowledgments

We thank Berk Tellioglu and Joseph Licata for their help with the experiments.

## References

Aasted, C. M., Yücel, M. A., Cooper, R. J., Dubb, J., Tsuzuki, D., Becerra, L., Petkov, M. P., Borsook, D., Dan, I., & Boas, D. A. (2015). Anatomical guidance for functional near-infrared spectroscopy: AtlasViewer tutorial. Neurophotonics, 2(2), 020801. 10.1117/1.NPh.2.2.020801PubMed

Aili, R., Zhou, S., Xu, X., He, X., & Lu, C. (2025). The cortical architecture representing the linguistic hierarchy of the conversational speech. NeuroImage, 311, 121180. 10.1016/j.neuroimage.2025.121180

Berni, T., Sacheli, L. M., Cicirello, M., Tettamanti, M., Paulesu, E., & Zapparoli, L. (2026). Together we sync: A systematic qualitative and quantitative review of fMRI hyper scanning studies. Social Cognitive and Affective Neuroscience, 21(1), saf122. 10.1093/scan/nsaf122 PubMed

Chen, D., Zhang, R., Lu, J., Wang, P., Bei, L., Lu, C., & Li, X. (2022). Gamma-band neural couplng durng conceptual algnment. Human Bran Mappng, 43(9), 2992–3006. 10.1002/hbm.25831

Czeszumsk, A., Liang, S. H.-Y., Dikker, S., König, P., Lee, C.-P., Koole, S. L., & Kelsen, B. (2022). Cooperatve behavor evokes nterbran synchrony n the prefrontal and temporoparetal cortex: A systematc revew and meta-analyss of fNIRS hyperscannng studes. eNeuro, 9(2), ENEURO.0268-21.2022. 10.1523/ENEURO.0268-21.2022eNeuro

De Felce, S., Chand, T., Croy, I., Engert, V., Goldstein, P., Holroyd, C. B., Kirsch, P., Krach, S., Ma, Y., Scheele, D., Schurz, M., Schweinberger, S. R., Hoehl, S., & Vrticka, P. (2025). Relatonal neuroscence: Insghts from hyperscannng research. Neuroscence & Bobehavoral Revews, 169, 105979. https://doi.org/10.1016/j.neubiorev.2024.105979 10.1016/j.neuborev.2024.105979

Delus, J. A. M., & Müller, V. (2023). Interpersonal synchrony when sngng n a chor. Frontersn Psychology, 13, 1087517. 10.3389/fpsyg.2022.1087517

Domínguez-Arrola, M. E., Lam, P. C. H., Pérez, A., & Pell, M. D. (2026). Interpersonal neural coordnaton tracks nteracton qualty durng naturalstc conversaton. NeuroImage, 338, 122090. 10.1016/j.neuromage.2026.122090 ScenceDrect

Ellngsen, D.-M., Isenburg, K., Jung, C., Lee, J., Gerber, J., Mawla, I., Sclocco, R., Grahl, A., Anzoln, A., Edwards, R. R., Kelley, J. M., Kirsch, I., Kaptchuk, T. J., & Napadow, V. (2023). Bran-to-bran mechansms underlyng pan empathy and socal modulaton of pan n the patent-clncan nteracton. Proceedngs of the Natonal Academy of Scences of the Unted States of Amerca, 120(26), e2212910120. 10.1073/pnas.2212910120 PubMed

Frth, C. D., & Frth, U. (2006). The neural bass of mentalzng. Neuron, 50(4), 531–534. 10.1016/j.neuron.2006.05.001

Gagnon, L., Yücel, M. A., Boas, D. A., & Cooper, R. J. (2014). Further mprovement n reducng superfcal contamnaton n NIRS usng double short separaton measurements. NeuroImage, 85, 127–135. 10.1016/j.neuromage.2013.01.073

Gan, R., Qiu, Y., Liao, J., Zhang, Y., Wu, J., Peng, X., Lee, T. M.-C., & Huang, R. (2024). Mappng the mentalzng bran: An ALE meta-analyss to dfferentate the representaton of socal scenes and ages on theory of mnd. Neuroscence & Bobehavoral Revews, 167, 105918. 10.1016/j.neuborev.2024.105918PubMed

Gvrts, H. Z., & Perlmutter, R. (2020). What guides us to neurally and behavorally algn wth anyone specfc? A neurobologcal model based on fNIRS hyperscannng studes. The Neuroscentst, 26(2), 108–116. 10.1177/1073858419861912

Healey, M. L., & Grossman, M. (2018). Cogntve and affectve perspectve-takng: Evdence for shared and dssocable anatomcal substrates. Fronters n Neurology, 9, 491. 10.3389/fneur.2018.00491

Hrsch, J., Noah, J. A., Zhang, X., Dravida, S., & Ono, Y. (2018). A cross-bran neural mechansm for human-to-human verbal communcaton. Socal Cogntve and Affectve Neuroscence, 13(9), 907–920. 10.1093/scan/nsy070

Hrsch, J., Zhang, X., Noah, J. A., & Ono, Y. (2017). Frontal temporal and paretal systems synchronze wthn and across brans durng lve eye-to-eye contact. NeuroImage, 157, 314–330. 10.1016/j.neuromage.2017.06.018

Holland, A. L. (1982). Observng functonal communcaton of aphasc adults. Journal of Speech and Hearng Dsorders, 47(1), 50–56. 10.1044/jshd.4701.50

Holmes, C. J., Hoge, R., Collins, L., Woods, R., Toga, A. W., & Evans, A. C. (1998). Enhancement of MR mages usng regstraton for sgnal averagng. Journal of Computer Asssted Tomography, 22(2), 324–333. 10.1097/00004728-199803000-00032 PubMed

Huppert, T. J., Diamond, S. G., Franceschn, M. A., & Boas, D. A. (2009). HomER: A revew of tme-seres analyss methods for near-nfrared spectroscopy of the bran. Appled Optcs, 48(10), D280–D298. 10.1364/AO.48.00D280 PubMed

şblr, E. E. (2016). Hyperscannng of bran-to-bran nteractonsn the prefrontal cortex durng a jont sentence readng task: An optcal bran magng study [Thess, Mddle East Techncal University].

Jang, J., Dai, B., Peng, D., Zhu, C., Lu, L., & Lu, C. (2012). Neural synchronzaton durng face-to-face communcaton. The Journal of Neuroscience, 32(45), 16064–16069. 10.1523/JNEUROSCI.2926-12.2012

Jang, L., Zou, Q., & Yu, S. (2026). EEG nter-bran synchrony n socal nteracton: A systematc revew and meta-analyss of condton effects and outcome assocatons. Bologcal Psychology, 210, 109367. 10.1016/j.bopsycho.2026.109367ResearchGate

Kawasak, M., Yamada, Y., Ushiku, Y., Myauch, E., & Yamaguchi, Y. (2013). Inter-bran synchronzaton durng coordnaton of speech rhythm n human-to-human socal nteracton. Scentfc Reports, 3, 1692. 10.1038/srep01692

Keller, P. E., Novembre, G., & Hove, M. J. (2014). Rhythm n jont acton: Psychologcal and neurophysologcal mechansms for real-tme nterpersonal coordnaton. Phlosophcal Transactons of the Royal Socety B: Bologcal Scences, 369(1658), 20130394. 10.1098/rstb.2013.0394

Kelsen, B. A., Sumich, A., Kasabov, N., Liang, S. H. Y., & Wang, G. Y. (2022). What has socal neuroscence learned from hyperscannng studes of spoken communcaton? A systematc revew. Neuroscence & Bobehavoral Revews, 132, 1249–1262. 10.1016/j.neuborev.2020.09.008

Knrech, S., Djalovski, A., Kraus, L., Louzoun, Y., & Feldman, R. (2017). Bran-to-bran synchrony durng naturalstc socal nteractons. Scentfc Reports, 7, 17060. 10.1038/s41598-017-17339-5

Lacade, C. M., Fulbright, R. K., Arora, J., Constable, R. T., & Papademetris, X. (2008). Brodmann areas defned n MNI space usng a new tracng tool n BoImage Sute. Organzaton for Human Bran Mappng Annual Meetng.

Lu, J., Zhang, R., Xie, E., Lin, Y., Chen, D., Lu, Y., Li, K., Chen, M., Li, Y., Wang, G., & L, X. (2023). Shared ntentonalty modulates nterpersonal neural synchronzaton at the establshment of communcaton system. Communcatons Bology, 6, 832. 10.1038/s42003-023-05197-z

Lu, Y., Piazza, E. A., Simony, E., Shewokis, P. A., Onaral, B., Hasson, U., & Ayaz, H. (2017). Measurng speaker-lstener neural couplng wth functonal near nfrared spectroscopy. Scentfc Reports, 7, 43293. 10.1038/srep43293

Lotter, L. D., Kohl, S. H., Gerloff, C., Bell, L., Niephaus, A., Kruppa, J. A., Dukart, J., Schulte-Rüther, M., Reindl, V., & Konrad, K. (2023). Revealng the neurobology underlyng nterpersonal neural synchronzaton wth multmodal data fuson. Neuroscence & Bobehavoral Revews, 146, 105042. 10.1016/j.neuborev.2023.105042

Merchant, J. S., Glaros, S., Edakoth, E., Harris, R., Tchangalova, N., & Redcay, E. (2025). Bran bases of real-tme socal nteracton: A meta-analytc nvestgaton of human neuromagng studes. Aperture Neuro, 5. 10.52294/001c.138339

Molav, B., & Dumont, G. A. (2012). Wavelet-based moton artfact removal for functonal nearnfrared spectroscopy. Physologcal Measurement, 33(2), 259–270. 10.1088/0967-3334/33/2/259

Ono, Y., Zhang, X., Noah, J. A., Dravida, S., & Hirsch, J. (2022). Bdrectonal connectvty between Broca’s area and Werncke’s area durng nteractve verbal communcaton. Bran Connectvty, 12(3), 210–222. 10.1089/bran.2020.0790

Osaka, N., Minamoto, T., Yaoi, K., Azuma, M., Shimada, Y. M., & Osaka, M. (2015). How two brans make one synchronzed mnd n the nferor frontal cortex: fNIRS-based hyperscannng durng cooperatve sngng. Fronters n Psychology, 6, 1811. 10.3389/fpsyg.2015.01811

Papademetrs, X., Jackowski, M. P., Rajeevan, N., DStaso, M., Okuda, H., Constable, R. T., & Stab, L. H. (2006). BoImage Sute: An ntegrated medcal mage analyss sute: An update. The Insght Journal, 2006, 209.

Pérez, A., Carreiras, M., & Duñabeta, J. A. (2017). Bran-to-bran entranment: EEG nterbran synchronzaton whle speakng and lstenng. Scentfc Reports, 7, 4190. 10.1038/s41598-017-04464-4

Pckerng, M. J., & Garrod, S. (2004). Toward a mechanstc psychology of dalogue. Behavoral and Bran Scences, 27(2), 169–225. 10.1017/S0140525X04000056 PubMed

Schlbach, L., Timmermans, B., Reddy, V., Costall, A., Bente, G., Schlicht, T., & Vogeley, K. (2013). Toward a second-person neuroscence. Behavoral and Bran Scences, 36(4), 393–414. 10.1017/S0140525X12000660PubMed

Schurz, M., Radua, J., Tholen, M. G., Maliske, L., Margulies, D. S., Mars, R. B., Sallet, J., & Kanske, P. (2021). Toward a herarchcal model of socal cognton: A neuromagng meta-analyss and ntegratve revew of empathy and theory of mnd. Psychologcal Bulletn, 147(3), 293–327. 10.1037/bul0000303

Shamay-Tsoory, S. G., Saporta, N., Marton-Alper, I. Z., & Gvirts, H. Z. (2019). Herdng brans: A core neural mechansm for socal algnment. Trends n Cogntve Scences, 23(3), 174–186. 10.1016/j.tcs.2019.01.002ScenceDrect

Speer, S. P. H., Mwlambwe-Tshlobo, L., Tsoi, L., Burns, S. M., Falk, E. B., & Tamir, D. I. (2024). Hyperscannng shows frends explore and strangers converge n conversaton. Nature Communcatons, 15, 7781. 10.1038/s41467-024-51990-7

Stephens, G. J., Silbert, L. J., & Hasson, U. (2010). Speaker-lstener neural couplng underles successful communcaton. Proceedngs of the Natonal Academy of Scences of the Unted States of Amerca, 107(32), 14425–14430. 10.1073/pnas.1008662107 PubMedCentral (PMC)

Stolk, A., Noordzij, M. L., Verhagen, L., Volman, I., Schoffelen, J.-M., Oostenveld, R., Hagoort, P., & Toni, I. (2014). Cerebral coherence between communicators marks the emergence of meanng. Proceedngs of the Natonal Academy of Scences of the Unted States of Amerca, 111(51), 18183–18188. 10.1073/pnas.1414886111

Van Overwalle, F. (2009). Socal cognton and the bran: A meta-analyss. Human Bran Mappng, 30(3), 829–858. 10.1002/hbm.20547

von Lühmann, A., Zheng, Y., Ortega-Martinez, A., Kiran, S., Somers, D. C., Cronn-Golomb, A., Awad, L. N., Ellis, T. D., Boas, D. A., & Yücel, M. A. (2021). Toward neuroscence of the everyday world (NEW) usng functonal near-nfrared spectroscopy. Current Opnon n Bomedcal Engneerng, 18, 100272. 10.1016/j.cobme.2021.100272 PubMed

Wang, X., Zhang, Y., He, Y., Lu, K., & Hao, N. (2022). Dynamc nter-bran networks correspond wth specfc communcaton behavors: Usng functonal near-nfrared spectroscopy hyperscannng durng creatve and non-creatve communcaton. Fronters n Human Neuroscence, 16, 907332. 10.3389/fnhum.2022.907332

Yücel, M. A., von Lühmann, A., Scholkmann, F., Gervain, J., Dan, I., Ayaz, H., Boas, D., Cooper, R. J., Culver, J., Elwell, C. E., Eggebrecht, A., Franceschn, M. A., Grova, C., Homae, F., Lesage, F., Obrig, H., Tachtsds, I., Tak, S., Tong, Y., … Wolf, M. (2021). Best practces for fNIRS publcatons. Neurophotoncs, 8(1), 012101. 10.1117/1.NPh.8.1.012101 PubMed

Zhao, Q., Zhao, W., Lu, C., Du, H., & Chi, P. (2024). Interpersonal neural synchronzaton durng socal nteractons n close relatonshps: A systematc revew and meta-analyss of fNIRS hyperscannng studes. Neuroscence & Bobehavoral Revews, 158, 105565. 10.1016/j.neuborev.2024.105565

